# Portable real-time colorimetric LAMP-device for rapid quantitative detection of nucleic acids in crude samples

**DOI:** 10.1101/2020.07.22.215251

**Authors:** G. Papadakis, A. K. Pantazis, N. Fikas, S. Chatziioannidou, V. Tsiakalou, K. Michaelidou, V. Pogka, M. Megariti, M. Vardaki, K. Giarentis, J. Heaney, E. Nastouli, T. Karamitros, A. Mentis, A. Zafiropoulos, G. Sourvinos, S. Agelaki, E. Gizeli

## Abstract

Loop-mediated isothermal amplification is known for its high sensitivity, specificity and tolerance to inhibiting-substances. We developed a device for performing real-time colorimetric LAMP combining the accuracy of lab-based quantitative molecular diagnosis with the simplicity of point-of-care testing. This handheld device employs a single reaction-pot for amplification and a mini-camera for detection. Competitive features are the rapid analysis (<30min), quantification over 9 log-units, crude sample-compatibility (saliva, tissue, swabs), low detection limit (<5copies/reaction), smartphone-operation and fast prototyping (3D-printing). The device’s clinical utility is demonstrated in cancer-mutations and COVID-19 testing. Excellent performance includes: detection of 0.01% of *BRAF*-V600E-to-wild-type molecules; 97% sensitivity to SARS-CoV-2 RNA detection (89 samples); 83% (Ct<34), 98% (Ct<30) and 100% (Ct<25) to 163 nasopharyngeal-swabs; 100% specificity in all cases. The device high technology-readiness-level makes it a suitable platform for performing any colorimetric LAMP assay; moreover, its simple and inexpensive fabrication holds promise for fast deployment and application in global diagnostics.

---

Over the past three decades, the invention of real-time quantitative PCR (qPCR) has brought a fundamental transformation in the field of molecular diagnostics^1-3^. Until today, the power of qPCR lies in the ability to quantify nucleic acids over an extraordinarily wide dynamic range and without the need for post-amplification manipulations^4^. However, qPCR is still a relatively complex, lab-based technology requiring thermal blocks with precise temperature control for amplification, expensive optical components for detection and sophisticated design of fluorescent probes for efficient DNA targeting or extensive optimization when using DNA dyes^5,6^. The development of alternative methodologies for operation outside a centralized lab and/or at the point-of-care (POC) in order to shorten lengthy procedures and provide fast results at a fraction of the cost is an on-going goal in the biotechnology and biomedical research communities.

Loop-mediated isothermal amplification (LAMP) stands out as a simpler yet powerful alternative to PCR of exceptional specificity in the case of sequence-specific amplification since it employs 4 to 6 primers which recognize 6 to 8 different regions in the target sequence^7^. LAMP can be performed at a constant temperature (60-65°C) eliminating the need for thermal cycling^8,9^, is faster^10, 11^ and less affected by inhibitors commonly present in biological samples^12, 13^. LAMP amplification can be monitored in real-time primarily through fluorescence or turbidimetry providing quantitative information^14-16^. The latter is important not only for lab-based but also for point-of-care applications as, for example, in the case of HIV viral load monitoring in resource-limited areas^14, 17^. However, the bulky and not so sensitive turbidimeters^18, 19^ and expensive real-time fluorescence-machines are not ideal for POC diagnostics. The same applies to the use of sophisticated but costly microfluidic chips^20, 21^ often used for real time monitoring in combination with optical^22^, electrochemical^23^, smartphone^24-26^, acoustic^27^ or other microdevices^22, 28, 29^. Moreover, fluorescent LAMP suffers similarly to PCR from high background and often inhibition of the reaction due to the increased dye concentration^30^.

As an alternative, researchers have been developing assays based on naked-eye observation^26, 31^ with end-point colorimetric LAMP being the most popular assay^32-37^. The above can employ simple means of heating e.g., a water bath, a thermos etc. or portable and/or battery operated instruments for isothermal amplification and eye-observation, all suitable for applications in resource-limited areas^34, 38^. Downsides of the naked-eye detection are the need of a “trained eye” for results interpretation, especially at low numbers of the target, and the qualitative nature of the results^26^. Attempts to overcome the above problem include the use of digital image analysis employed in microfluidic chips with nanoliter volumes for end-point color quantification^39^, a sophisticated but rather complex method. In another work, a photonic crystal-setup was employed to achieve naked-eye quantification of LAMP reactions after several rinsing and drying steps and exposure of LAMP amplicons to the surrounding space^40^. Finally, LAMP products combined with a lateral flow dipstick have been used for highly specific and visual detection of amplicons at the POC^26^. The paper-based strips are competitive end-point LAMP candidates for lab-free testing since they do not require any instrumentation; however, they are qualitative and prone to contamination^26^. As concluded in a recent review^7^, while open fluorescence-based platforms are already commercially available and can be implemented for lab-based LAMP detection, tailored-made devices’ adaptation to operational environments is still missing.

In this work, we report the design and construction of a portable biomedical device for performing real-time quantitative colorimetric LAMP (qcLAMP) for a broad range of applications in a lab or POC setting. One of the main advantages of the method is the use of a single reaction-pot (Eppendorf-tube) for simultaneous amplification and quantitative detection obviating completely the need for a microfluidic chip and reducing the risk for contamination. A schematic of the qcLAMP device and concept is presented in Fig. 1a. The device is constructed using miniaturized electronic components produced by three-dimensional (3D) additive manufacturing and operates via an in-house developed smartphone application. For monitoring the color change during DNA amplification, a novel way of heating was introduced allowing efficient amplification with parallel visualization of the reaction by a mini digital camera controlled by a Raspberry Pi. The above, when combined with an application for digital image analysis, can extract rapidly quantitative information for a wide dynamic range of the genetic target. For the proof-of-concept, we used the qcLAMP device for the detection of SARS-CoV-2 RNA, extracted from patients’ samples or directly from nasopharengeal (NP) specimens. We also used the detection of the *BRAF* V600E mutation from genomic DNA derived from formalin-fixed paraffin-embedded (FFPE) samples as an example of companion diagnostics and pharmacogenomics. Our results prove both the suitability of the newly reported real-time qcLAMP as a reliable and versatile method for genetic testing as well as the device high technology readiness level allowing immediate implementation in several applications and settings.

**Figure 1:**
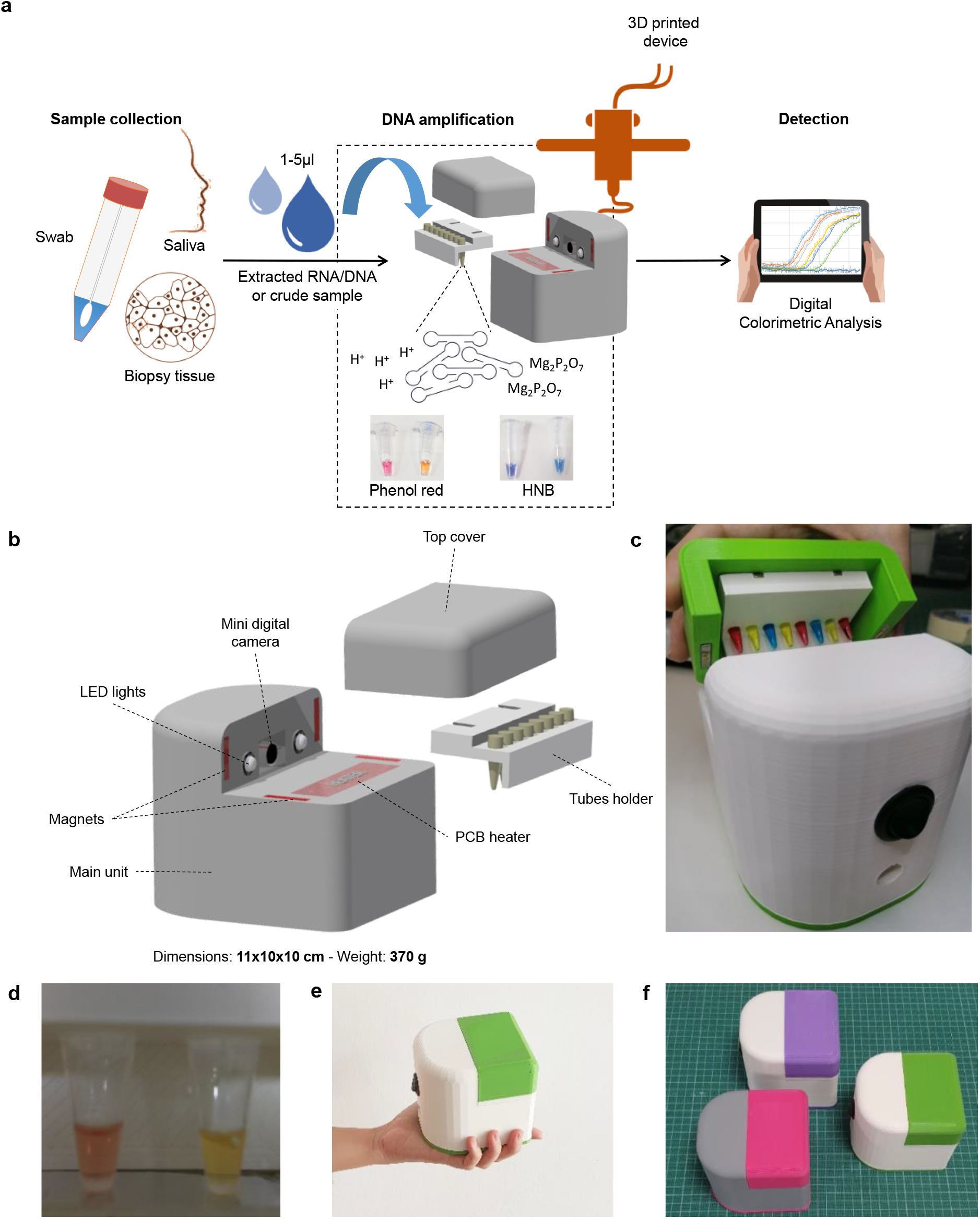
(a) Overview of the real-time quantitative colorimetric LAMP (qcLAMP) concept. Bottom: (b) Schematic representation of the qcLAMP device and components. (c) Photograph of the device and tubes-holder. (d) Captured image of two Eppendorf-tubes inside the chamber and during operation of the device. (e) Image of the handheld device for performing qcLAMP. (f) Three prototype devices of various colors; parallel production of two boxes requires ∼12 h.

## Results

### Device design and construction

The real-time colorimetric device was fabricated using additive manufacturing. Following the design of the device main housing unit, digital manufacturing based on 3D-printing was employed. Within this unit, all the required electronic components and an 8-slot tube-holder accommodating standard 0.2 ml Eppendorf tubes were positioned according to a CAD design; moreover, a top cover was used to apply pressure on the tubes and simultaneously provide an isolated chamber from the environment while real-time colorimetric LAMP detection takes place (Fig. 1b and c). The electronic components consist of a main electronic board, a Raspberry Pi Zero, a Raspberry Zero camera and a temperature sensor connected to a PCB resistive microheater. When the top part is positioned in place for operation, the mini camera is aligned opposite the tubes while two led lights are placed on both sides to provide stable lighting conditions in the chamber (Fig. 1d). The heating element is in direct contact only with the bottom of the tubes (∼2 mm diameter of contact surface), while application of appropriate pressure (∼2 MPa) from the top cover maintains good contact with the heating element. This particular setup allows for the direct inspection of the reaction progress through the side of the tube-walls combined with efficient and timely heating of the solution. The tube holder and cover top parts are held in place with the aid of magnets, which facilitate the opening and closing of the device and eliminate the need of screws. Additionally, the magnets minimize any potential variation in the applied pressure among different runs of the device.

The dimensions (11×10×10 cm) and weight (370 g) of the device are ideal for use at the POC (Fig. 1e). The housing unit is made mostly of polylactic acid (PLA) plastic and requires ∼12 hours of continuous printing while the assembly of all the components can be completed in 20 min. The simple construction requirements allow for the production of several prototypes in less than a week (Fig. 1f). The device connects via Bluetooth to a smartphone or tablet and operates through an in-house developed Android application (Fig. S1).

### Digital image analysis and performance evaluation

Colorimetric LAMP detection typically relies on the naked-eye evaluation of color change which is achieved through the use of different indicators such as pH^41^, metal binding^42^ or DNA binding^43^ dyes. Since color change can often be difficult to discern under variable conditions (lighting, target concentration, etc.), smartphone cameras combined with algorithms or radiometric imaging have been proposed as a generic solution for more accurate and quantitative colorimetric test –results^39, 43-48^.

Here, a mini digital camera is employed for monitoring in real-time the transition through various color shades during colorimetric LAMP amplification. The camera collects non-calibrated images at predefined time intervals (6 sec minimum interval) and automatically extracts the red, green and blue (RGB) pixel values. A real-time curve is displayed plotting the difference between green and blue or green and red pixels, depending on the indicator dye. The selection of these two formulas was derived after systematic analysis of a series of images containing either the phenol red or the hydroxy napthol blue (HNB) dyes (Fig. S2a-e). When the phenol red indicator was used, the Green-Blue pixel formula responded well into discriminating the positive from the negative sample; with the HNB indicator, the Green-Red formula resulted in better discrimination.

Based on the above, we programmed the qcLAMP device so that the end-user could select any of the two formulas. The formulas are displayed in the Y-axis of a real-time plot as color index units (pixels). Figures 2a and 2b show examples of real time colorimetric LAMP reactions performed in the device at the phenol red and the HNB channels, respectively. Whole bacteria cells were used as template for the positive reactions which were run in parallel with negative control reactions without template. Regardless of the dye used in each reaction, the change in the slope of the line corresponding to the positive sample at a specific time point (14.1 and 13.3 min for the phenol red and HNB, respectively) indicates the presence of the target, as opposed to the line corresponding to the negative sample which remains almost flat throughout the monitoring period.

**Figure 2:**
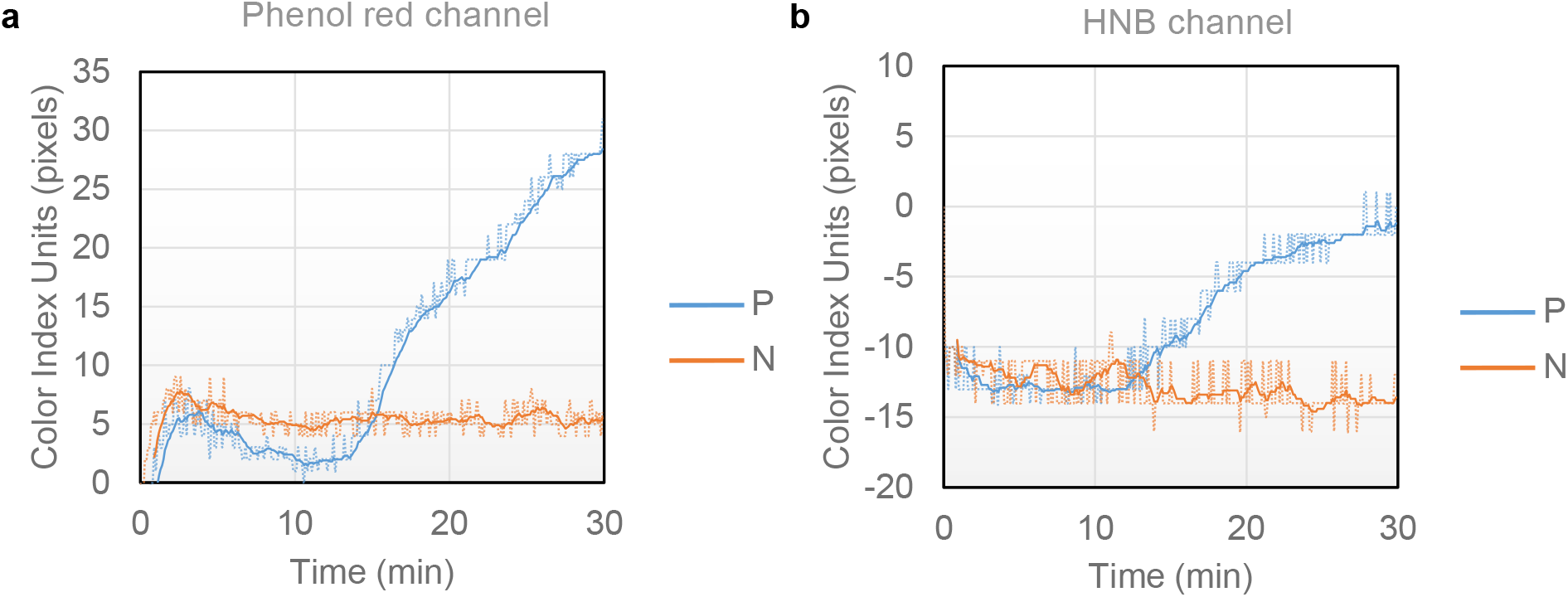
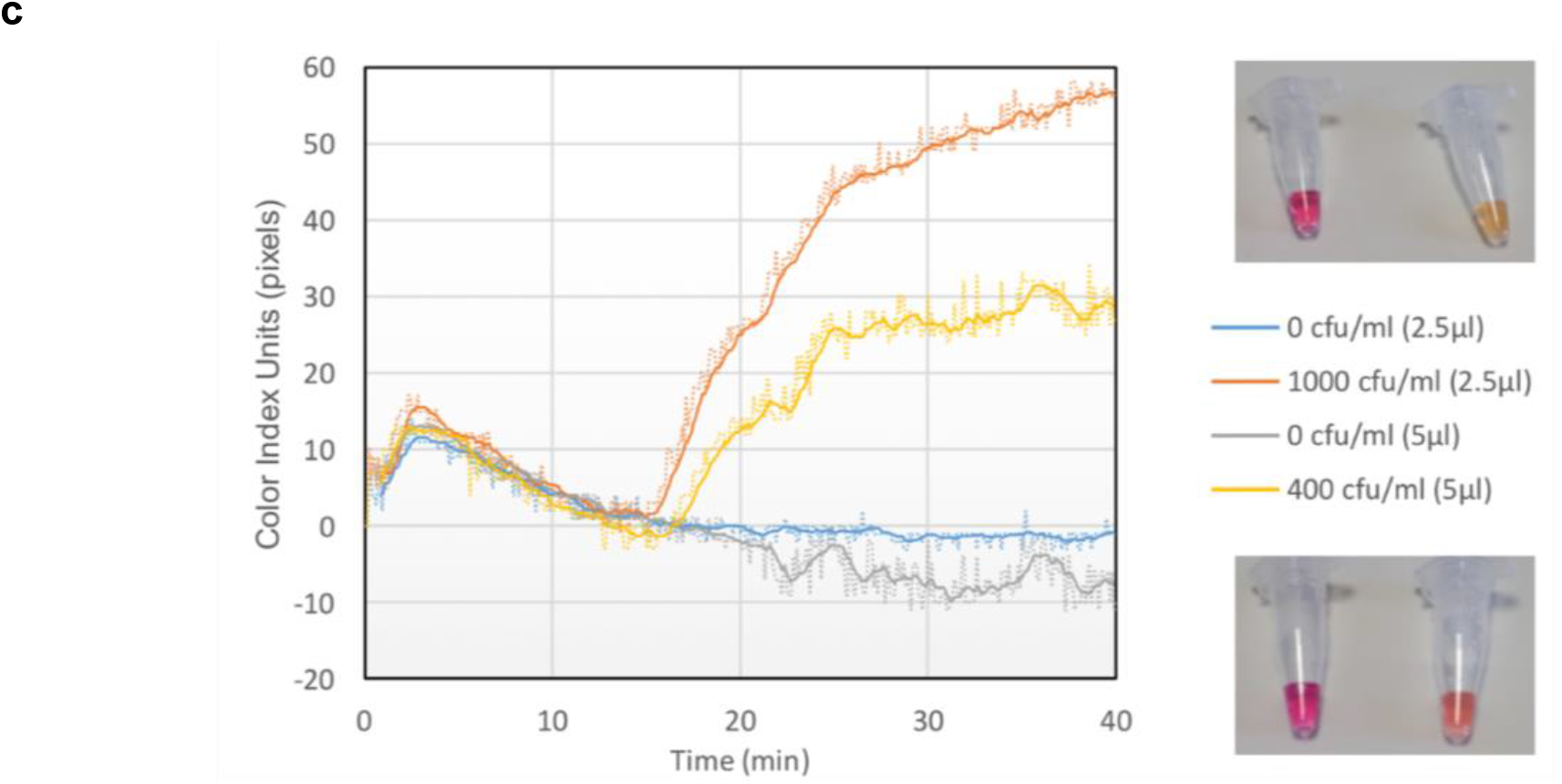
(a) Real-time colorimetric LAMP reactions performed inside the device using the phenol red indicator. The positive reaction (blue line) contained 10 lysed bacteria as template while the negative control contained no bacteria (orange line). (b) Similar reactions to (a), this time performed with the HNB indicator. In both cases, the heating process started at t=0 min and the heater reached the required temperature (63°C) after approximately 2 min. All reactions were stopped after 30 minutes. (c) Left: Real time colorimetric curves monitored during qcLAMP amplification performed with 2 different bacteria concentrations (400 and 1000 CFU/ml) spiked in saliva samples. The two concentrations were chosen based on previously reported detection limits of bacteria using end-point colorimetric LAMP or biosensors (*13, 54*). Both were successfully detected in less than 17 min while zero background signal was monitored for the negative controls. Right, top: Picture of end-point reactions with 0 and 1000 CFU/ml after 40 min of incubation. Right bottom: Picture of end-point reactions with 0 and 400 CFU/ml after 40 min of incubation.

For the performance evaluation of the qcLAMP, we investigated the mechanical robustness of the device as well as the speed, efficiency and sensitivity during LAMP amplification of a genetic target (*Salmonella InvA* gene). Regarding the former, potential variations in the measurements introduced by the position of the tube inside the holder were tested. Results showed excellent reproducibility in terms of the average time at which the color index slope starts to increase, i.e., a 24 sec deviation (15.7±0.4 min, i.e. 2.5%) when phenol read indicator was used and 48 sec (17.9±0.8 min, i.e. 4.5%) when HNB was applied (Fig. S3a). This is a small variation taking into consideration that the reactions were different preparations and not from a single master mix. Regarding the reaction-speed, the difference between qcLAMP and real-time fluorescent LAMP detection was found to be around 5.5 min (Fig. S3b); this difference is even smaller (by approximately 2 min) if we consider the fact that real-time fluorescent LAMP measurements start only after the heat block has reached the required temperature (63°C) while in the qcLAMP device heating starts at t=0 min when the heater is still at room temperature. Finally, the efficiency and sensitivity of qcLAMP inside a complex sample was initially demonstrated in saliva, during the detection of down to 0.4 CFU/μl (Fig. 2c). The use of 5 μl of crude saliva in a 25 μl LAMP reaction translates to a limit of detection of 2 CFUs per reaction (0.08 copies/μl) and an impressive final crude sample concentration of 20% in contrast to the commonly required 10% or less.

Overall, the device showed excellent stability and reproducibility during the continuous and undisturbed operation of a testing period of over 12 months; using a 10000 mAh power-bank, the device was able to run non-stop for approximately 5 h (Fig. S4).

### Quantitative detection of *BRAF* V600E mutation from FFPE tissues and clinical validation

To validate our methodology for clinical applications, first we employed the qcLAMP device to detect various ratios of the *BRAF* V600E mutation in the presence of wild type (wt) alleles, a test of significance to pharmacogenomics and companion diagnostics. Isolated genomic DNA (from patient samples) containing the mutation of interest (mut) was mixed with wild type DNA (wt) to a total amount of 100 ng DNA in the following (mut:wt) ratios: 50%-10%-5%-1%-0.1%-0.01%-0%. The lowest ratio (0.01%), corresponding to the limit of detection, contained 10 pg of mutant DNA or approximately 2-3 mutant copies^49^. Triplicate LAMP reactions were measured in random order in the phenol red channel for 45 min in total. All different ratios showed a positive response within the first 30 min while for the 0% (or 100 ng wt) no signal change was observed. Figure 3a illustrates a set of real-time colorimetric curves for the whole range of measured ratios. The average time response for each ratio showed excellent correlation (R^2^=0.99) with the starting template concentration of mutant DNA (Fig. 3b). Overall, Fig. 3b demonstrates the ability of the method to provide positive results in a wide dynamic range of more than four orders of magnitude (10 pg to 50 ng of mutant genomic DNA) and with zero background signal when only wild type DNA is used. The requirement of 100 ng genomic DNA per reaction is an amount readily obtained from one 5-μm tissue section and similar to what other methods require^50, 51^.

**Figure 3:**
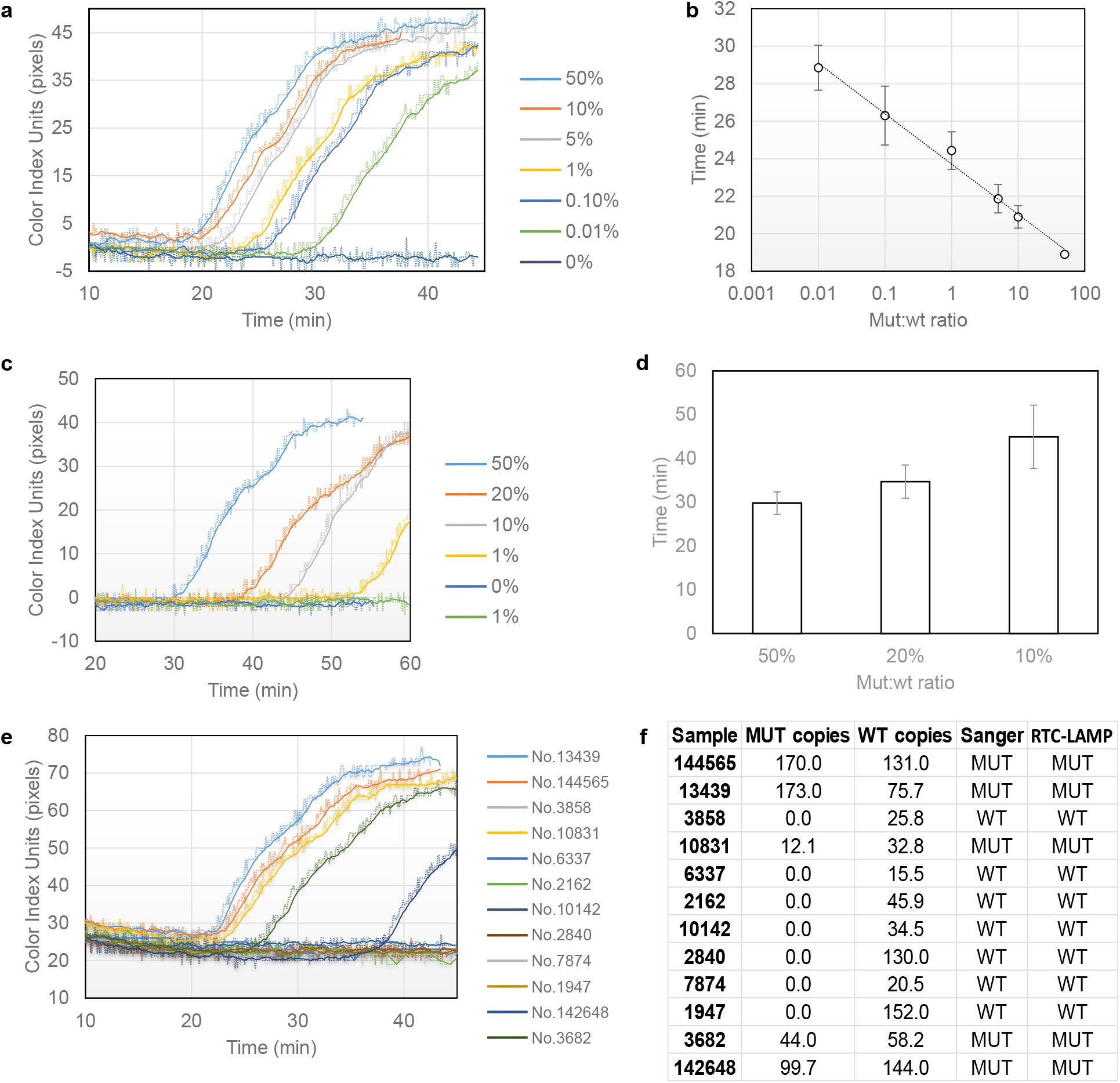
(a) Real-time colorimetric curves corresponding to mut:wt starting template ratios spanning more than four orders of magnitude. (b) Calibration curve produced from real-time data of triplicate colorimetric LAMP measurements with error bars representing standard deviation; y = −1.165ln(x) + 23.72 (R^2^ = 0.99). (c) Real-time data corresponding to tissue samples of various mut:wt ratios. The 1% was not consistently produced (yellow vs green line). (d) Comparison of real-time data for 10, 20 and 50% mut:wt tissue samples measured directly in the qcLAMPdevice without DNA purification. (e) Real time colorimetric LAMP results of 12 patient samples analyzed with the qcLAMP 3D device (blind tests). (f) Comparison of ddPCR, Sanger and qcLAMP test results of 12 clinical samples. The ddPCR method provided additional information regarding the concentration of mutant and wild type copies per analyzed sample.

Moreover, we investigated the possibility of performing direct detection of the *BRAF* V600E mutation from FFPE tissue without DNA extraction. Slices of paraffin tissue were collected and analyzed as described in the methods section; 7.5-10 μl of the solution were then transferred into LAMP reactions containing the phenol red indicator and placed in the 3D-device for 60 min. As expected, the wt tissue sample did not show any signal change. Figure 3c illustrates a set of real-time colorimetric curves for the 5 different ratios of mut:wt tissue samples (50-20-10-1-0%). The 50% tissue samples were found positive in approximately 30 min while the 20 and 10% after 35 and 45 min, respectively (Fig. 3d). Mixing wt and 10% preparations to a final ratio of 1% mut:wt was also found positive at 45.5 min on average but the result was reproduced only in 2 out of the 8 attempts.

The detection of the mutation in the 1% ratio was not always possible for two main reasons. Firstly, the LAMP reaction contained a fraction (10 μl) of a DNA preparation (1/2 of a 15-μm tissue slice melted in 100 μl water) that was not expected to contain more than 20 ng of genomic DNA. Secondly, the DNA preparation through melting was of low purity compared to standard DNA extraction methods that utilize protein digestion and overnight lysis steps^52, 53^. Larger slices of tissue, not available in the current study, could improve the reproducibility for the 1% ratio. The melting process could also be improved by using a combination of a commercially available lysis solution with a DNA intercalating colorimetric dye (e.g. crystal violet). In this study, melting in water was employed not only as a simple and cost-effective solution but also for compatibility issues with the pH dependent phenol red color indicator.

Finally, the presence of the *BRAF* V600E mutation in genomic DNA extracted from 12 clinical biopsy samples from cancer patients was investigated in blind tests with our device. A sample of 100 ng of extracted genomic DNA were tested in the phenol red channel while the same DNA preparation was used for Sanger sequencing and the ddPCR method. The presence of the mutation was verified by an increase in the color index units in five out of the 12 tested samples while the rest showed no change in pixels (Fig. 3e). Four of the positive samples were identified in less than 30 and one in less than 40 min. These results were in total agreement with Sanger sequencing method and the droplet digital PCR (ddPCR). According to ddPCR, the concentration of mutant copies successfully identified by the device ranged from 12.1 to 173 copies/μl (Fig. 3f).

### qcLAMP testing for SARS-CoV-2

In a second example, we validated the qcLAMP for infectious disease-testing focusing on the recent COVID-19 pandemic and urgent need for SARS-CoV-2 detection. Initially, the capability of qcLAMP to provide quantitative results was tested by deriving a calibration curve from a wide range of starting template material. For this curve, DNA derived from Influenza A was used as a reference molecule within the range of 10^1^ to 10^9^ copies using a 10-fold dilution. Figure 4a shows a calibration curve produced by triplicate measurements for Influenza A detection (Fig. S5a). The highest template concentration showed a time-to-positive result at 11.8±0.2 min while the lowest at 20.0±2.4 min. The achieved detection limit was 10 copies per reaction or 0.4 copies per μl input. The reproducibility of our device in respect to the same initial target concentration (10^9^ copies/reaction) was evaluated by monitoring 21 positive and 7 negative LAMP reactions. The standard deviation for the average time-to-positive results was 36 sec in an average response time of 10.2 min (Fig. S5b); no increase in the color index units was detected with the negative reactions (Fig. S5c).

**Figure 4:**
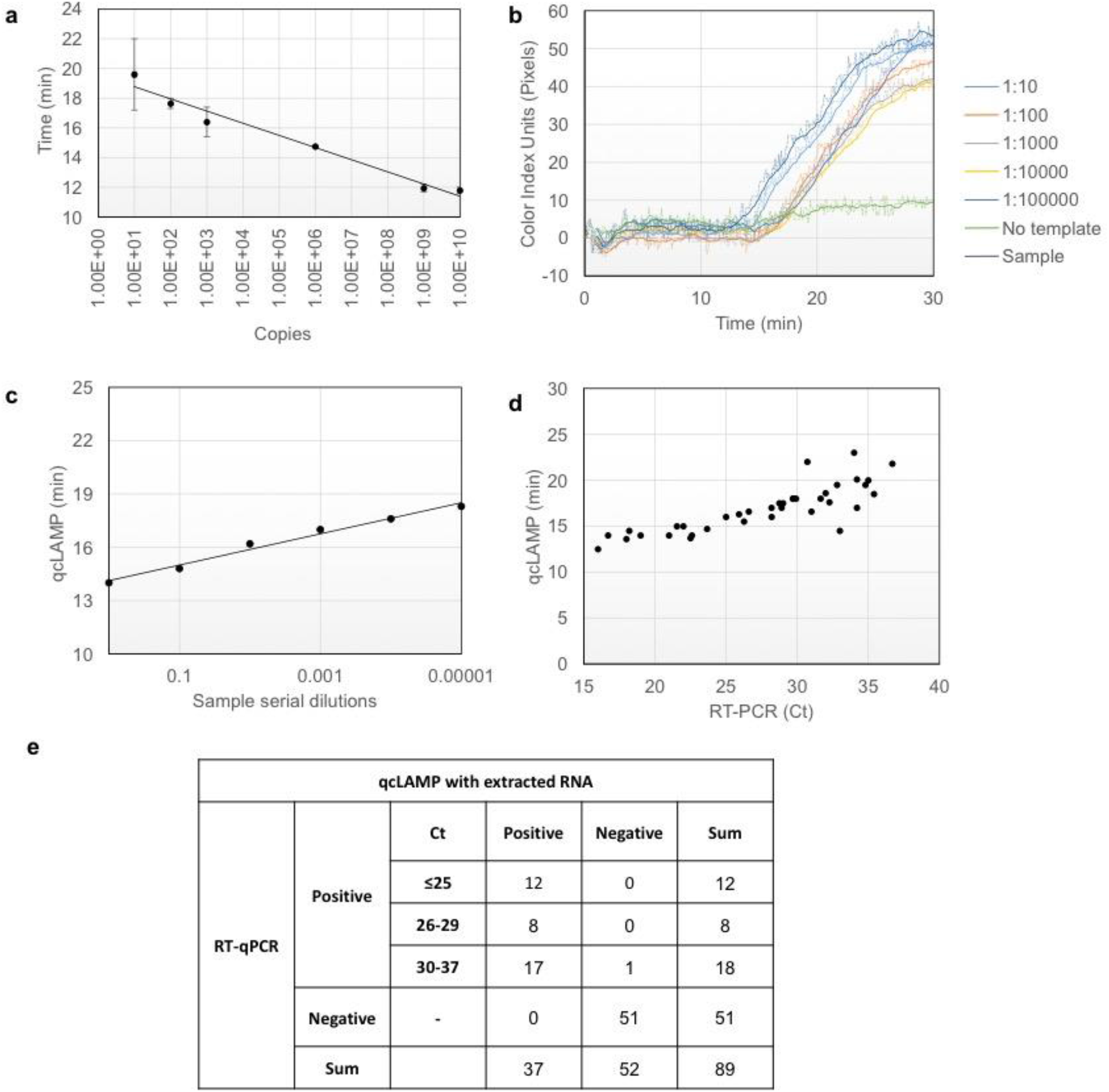
(a) Calibration curve using Influenza A DNA template ranging from 10^1^ to 10^9^ copies per reaction; error bars represent standard deviation of at least triplicates (b) Serial dilutions of a SARS-COV-2 positive sample with a reported Ct value of 19 were measured with the qcLAMP method. (c) Correlation (R^2^=0.99) between the qcLAMP time-to-positive results and the viral RNA concentration based on a 10-fold serial dilutions. (d) Scatter plot of the Ct values (ranging from 16 to ∼37) for 38 positive samples versus the qcLAMP time-to-positive (ranging from 12.5 to 23 min). qcLAMP measurements were performed in different days within 3 weeks using stored (−80°C) RNA samples. Only one point was missed by qCLAMP in the first run corresponding to a Ct of 35. (e) Diagnostic results of SARS-CoV-2 qcLAMP assay-evaluation with 89 clinical samples including 38 COVID-19 positive and 51 negative. An overall sensitivity of 97% (95% CI: 93-100), specificity of 100% and negative likelihood ratio of 0.026 (95% CI: 0.004-0.182) were calculated.

We then evaluated the performance of the device for the detection of SARS-CoV-2 during simultaneous reverse transcription and isothermal amplification targeting the *N* gene in clinical samples, starting from RNA extracted from patients’ nasopharyngeal or oropharyngeal swabs. We initially performed a set of qcLAMP measurements with serial dilutions of extracted total RNA from a SARS-CoV-2 positive sample with a reported Ct value of 19 (Fig. 4b). The real-time colorimetric curves were used to correlate the time-to-positive results with the viral RNA concentrations (Fig. 4c). A very good correlation was observed (R^2^=0.99) which verified the ability to extract quantitative information using the newly developed method. Furthermore, by employing a commercially available kit we managed to detect down to 5 copies of viral RNA per LAMP reaction within approximately 20 min (Fig. S5d).

Additionally, 89 patient samples consisting of 38 qRT-PCR positive and 51 negative for SARS-CoV-2 were tested with the qcLAMP method. The positive samples had reported qRT-PCR Ct values ranging from 16 to 36.8. The fastest observed time-to-positive result with qcLAMP was 12.5 min for a sample with a Ct value of 16 while the maximum was 23 min for a Ct of 36.7. Thirty-seven (37/38) of the positive samples (with respect to qRT-PCR) were successfully identified with qcLAMP in the first attempt. Only one (1/38) was identified as positive after a second repeat on a second run. The correlation of qRT-PCR Ct values and time-to-positive results with qcLAMP is shown in Fig. 4d. It should be noted that these were independent runs performed in different days within 3 weeks. In parallel to *N* gene-testing, detection of the human RNA target (*RNase* P) was carried out as a positive control.

Forty-seven (47/51) negative samples for SARS-CoV-2, 6 of which had been found positive for other viruses (Influenza, CMV, RSV, Adenovirus, and Enterovirus) were successfully identified as negative without any indication of cross-reactivity. Furthermore, we found that four (4/51) of the stored samples previously identified as SARS-CoV-2 positive were negative with our device. Running a second qRT-PCR verified our results; this observation was attributed to poor RNA quality after freezing and thawing. Finally, among the 89 samples used, 6 positive and 6 negative were blindly tested and identified with our method with 100% accuracy. The results from this study have important implications in the context of developing reliable molecular diagnostic tools, given the demonstrated excellent performance of the qcLAMP methodology with 0% of false positive and only 2.6% false negative results (Fig. 4e).

In a follow-up study, we assessed the capability of the qcLAMP device for the point-of-care, i.e., the detection of SARS-CoV-2 directly in crude samples. Currently, for SARS-CoV-2 collection the nasal / oropharyngeal swab sample is placed inside 1-3 ml of viral transport medium (VTM). Different types of commercial VTM samples can include various buffers but also dyes, some of which can inhibit the colorimetric assay and/or reduce the efficiency of the qcLAMP to discriminate between a positive and negative sample. For this reason, we performed a range of studies for the development of an optimized assay starting directly from a VTM-preserved nasopharyngeal specimen. Optimization parameters included the type of the transport medium used, dialysis buffer, sample dilution-volume and colorimetric dye. We selected to study clinical samples inside three commercial viral transport media (VTM) (Citoswab, Improviral, Liofilchem) diluted with a neutralizing solution. Dilutions of 1:1 were prepared and placed in the qcLAMp device for *N* gene amplification (63°C, 30 min) and colorimetric real-time detection (Fig. 5a). For validation, 163 nasopharyngeal patients-samples were selected randomly for specimens collected for RT-PCR testing, including both frozen (111) and fresh (52) ones. Of the 96 PCR-reported positive samples, we accurately identified 80 of them, with a time-to-positive result varying from 14.8 (Ct=8) to maximum 28.5 min (Ct=34), respectively (Fig. 5b). Samples with a Ct>30 were normally tested twice using aliquots withdrawn at different times and within 24 h. According to our results (Fig. 5b and 5c), no significant difference is observed in the qcLAMP detection capability between fresh and frozen specimens. In addition, all PCR-confirmed negative to SARS-Cov-2 samples (67/67) were correctly identified as negative with the qcLAMP method (Fig. 5c and 5d). When taking into account all our results (Ct<34), we calculated a sensitivity of 83% (95% CI: 77-92), specificity of 100% and a negative likelihood ratio of 0.17 (95% CI: 0.11-0.26).

**Figure 5:**
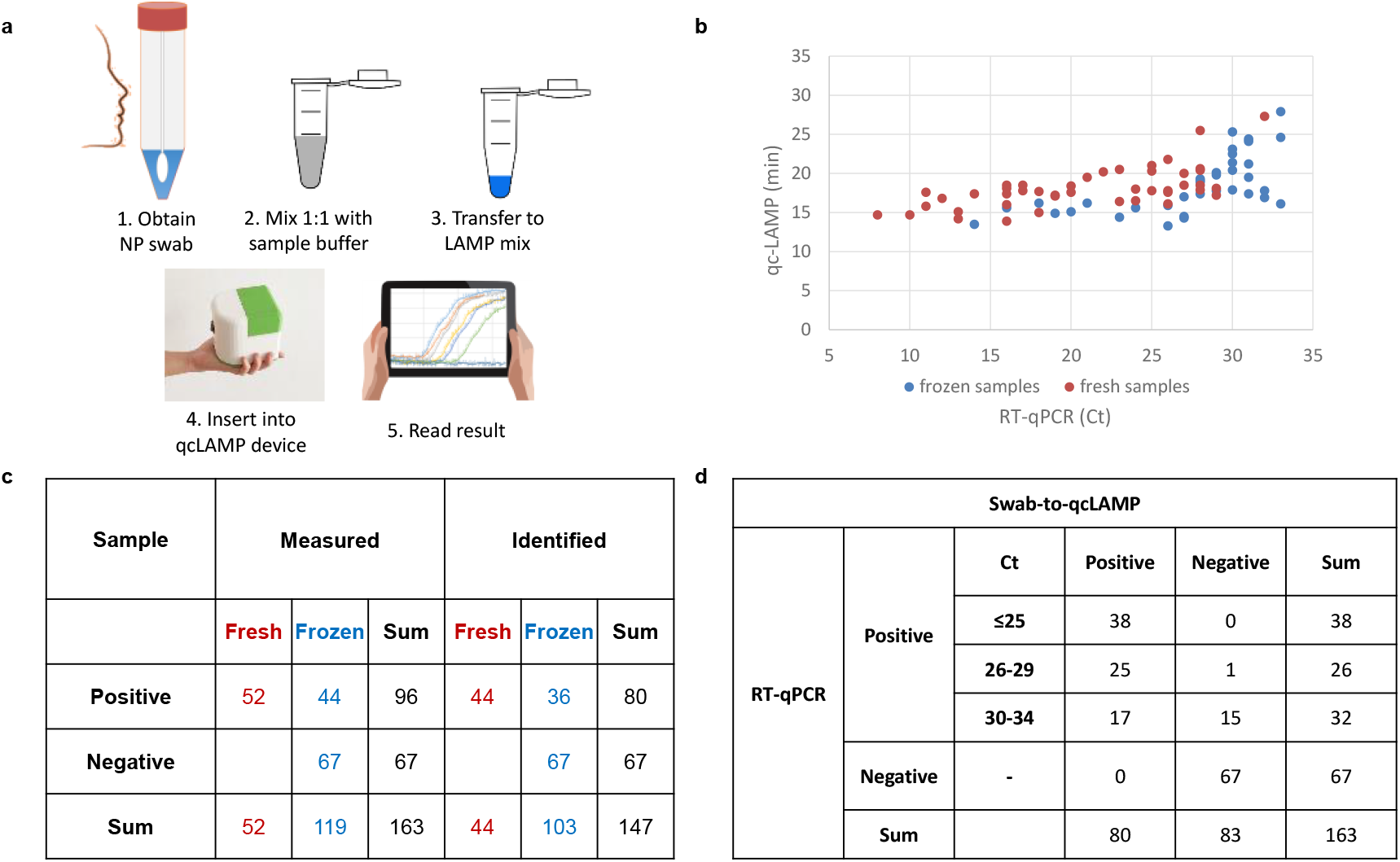
(a) Schematic representation of the workflow of the SARS-CoV-2 swab-to-qcLAMP detection, i.e. without RNA purification. (b) Scattered plot of the Ct values of PCR-confirmed positive samples by using the HNB colorimetric dye inside the LAMP-mix. (c) Table presenting overall diagnostic sensitivity and specificity of the qcLAMP for fresh and frozen samples; the red and blue colors are used to guide the eye and depict fresh and frozen samples, respectively. (d) when considering various Ct cut off values. For Ct<30 sensitivity becomes 98% (95% CI: 95-101) and negative likelihood ratio 0.016 (95% CI: 0.002-0.109); for Ct<25, sensitivity is calculated to be 100%.

## Discussion

Low-cost portable and quantitative methods for nucleic acid detection are desirable in molecular diagnostics both for lab and POC settings. Colorimetric LAMP, one of the simplest and most suitable formats for such applications is based primarily on end-point measurements deriving qualitative results. Here we describe a device that offers for the first time the ability to perform real-time colorimetric LAMP inside a single reaction-pot through an embedded mini-camera and extract quantitative information in a similar way to real-time fluorescent nucleic acid amplification methods but without their associated complexity and high cost. The newly developed device is based on a contamination-free closed tube format obviating the need for microfluidic chips, peristaltic pumps, fluorescent detectors and multiple step protocols employed elsewhere ^13, 26, 39, 40, 47, 48, 54^. It can achieve a similar limit of detection to the real-time fluorescent PCR and LAMP protocols (<5 copies/reaction, i.e., <0.2 copies/μl) while it does not suffer from background non-specific signal or need for signal drift correction or precise selection of threshold values required in other more sophisticated systems^27, 28^. It displays similar time-to-result with real-time fluorescent LAMP (Fig. S3); in addition, it can be performed in the presence of crude samples, such as 20% saliva (Fig. S4), tissue-specimens and nasopharyngeal swabs without prior nucleic acid extraction.

The clinical utility and broad applicability of the new technology is demonstrated in two applications. First, the qcLAMP device was applied as a companion diagnostic tool for *BRAF* V600E mutation-detection. The demonstrated ability to detect as low as 2-3 mutant copies within an abundance of wild-type background alleles (0.01% ratio) was more than two orders of magnitude better than commonly employed methods (Table 1). Among them are the Sanger sequencing which is the clinical standard, pyrosequencing, the Cobas test and the Infinity assay, all of which have a reported LOD of ≥5%^50, 55, 56^. Only the ddPCR has a better reported LOD of 0.001%^51^; however, ddPCR is expensive and unsuitable for companion diagnostics. Moreover, all 12 clinical samples tested blindly were identified according to Sanger sequencing and ddPCR. The current application suggests the potential of qcLAMP as a valuable tool for on-site companion diagnostic tests, when medical treatment is required, alleviating delays associated with testing in centralized labs ^57^. Such examples include the prescription of vemurafenib to melanoma patients carrying the *BRAF* V600E or warfarin and clopidogrel (Plavix) drugs prescribed for cardiovascular disease-treatment, the response of which is affected by genetic polymorphisms (*VKORC1, CYP2C9* and *CYP2C19* genes).

**Table 1:** Comparison of the qcLAMP, Sanger sequencing, ddPCR and qRT-PCR methods for the two applications. PCR^1^ and PCR^2^ refer to the different protocols used in the two validation sites (Lab. of Clinical Virology and Hellenic Pasteur Institute, respectively).

| **BRAF V600E ASSAY** |  |  |  | **SARS-CoV-2 ASSAY** |  |  |  |
| --- | --- | --- | --- | --- | --- | --- | --- |
|  | **qcLAMP** | **Sanger** | **ddPCR** | **qcLAMP<sup>crude</sup>** | **qRT-PCR<sup>1</sup>** | **qcLAMP** | **qRT-PCR<sup>2</sup>** |
| **Steps** | 1 | 2 | 2 | 1 | 2 | 1 | 2 |
| **Temperature** | 65°C | multiple | multiple | 65°C | 53°C (RT)<br>(60°C–95°C)<br>x40 cycles | 65°C | 50°C (RT)<br>(55°C–95°C)<br>x45 cycles |
| **Assay time** | N/A | N/A | N/A | 30 min | 90 min | 30 min | 80 min |
| **Sample-to-result time** | <2h | >24h | >2h | <40 min<br>POC | >3 h<br>lab-based | 2 h<br>lab-based | >3 h<br>lab-based |
| **Target/control** | *BRAF*<br>V600E/WT | *BRAF*<br>V600E/WT | *BRAF*<br>V600E/WT | *N* gene/<br>*RNase P* | *ORF/S/N*<br>gene/Internal<br>control | *N* gene/<br>*RNase P* | *RdRp* gene/<br>*RNase P* |
| **Preparatory steps** | 1 (DNA<br>extraction<br>or tissue<br>heating) | 2 (DNA<br>extraction &<br>thermal<br>cycling) | 2 (DNA<br>extraction &<br>droplets<br>formation) | Crude N/O-P<br>sample | NA<br>extraction | NA<br>extraction | NA<br>extraction |
| **Analytical performance** | 0.01%<br>(mut:wt<br>ratio) | 5%<br>(mut:wt<br>ratio) | 0.001%<br>(mut:wt ratio) | Not available | 1 copy/<br>reaction | 5 copies/<br>reaction | Not available |

The qcLAMP methodology was further used for the development of an assay for SARS-CoV-2 RNA detection and validation using COVID19 patients’ samples and against the standard qRT-PCR method recommended by the WHO and the CDC. The targeted *N* gene, selected for the proof-of-concept, was shown to be specific for SARS-CoV-2 since samples positive for respiratory or other viruses exhibited no cross-reactivity. Compared to qRT-PCR, the reverse transcription-qcLAMP method using extracted RNA delivers results within 23 min demonstrating at the same time very good sensitivity (97% for Ct<37) and specificity (100%) (Table 1). High throughput screening using centralized PCR cannot be achieved with the current qcLAMP device, since the latter can only process eight samples at a time. However, combined with a rapid extraction method, the qcLAMP can still have an impact in a minimally-equipped environment when urgent rapid decision-making is necessary, e.g. deciding on therapy options in the intensive care unit or screening of patients during scheduled (surgery) or unscheduled (maternity units) admissions, etc.

For the full exploitation of the qcLAMP methodology, we also developed a swab-to-qcLAMP assay combined with reverse-transcription, circumventing the RNA-extraction step and, thus, suitable for POC testing. Our results indicate an excellent specificity (100%) but reduced sensitivity (83% for Ct<34) when compared to the RNA-extracted assay, in agreement with previous studies^58-60^. By employing a Ct cut-off of <30 and <25, we calculated a sensitivity of 98% (64 samples) and 100% (38 samples), respectively. To evaluate the potential of the qcLAMP for POC testing, we compared our results to similar studies including validation of RT-LAMP assays with a comparable number of unprocessed naso/ortho-pharyngeal samples of a known Ct. To the best of our knowledge, the performance of the qcLAMP is better in terms of sensitivity and similar in terms of detection time with all LAMP-based SARS-Cov-2 assays reported so far, using naked eye end-point colorimetric^61, 62^, real-time spectrophotometric^59^ or fluorescent^58, 61, 63^ detection.

Our results complement current efforts by the scientific community to provide a reliable platform for nucleic acids-testing at the POC as a response to the emerging COVID-19 pandemic. Notable examples are first, an accessible and portable POC instrument for RT-LAMP detection of SARS-CoV-2 using unpurified NP swabs^64^. This work also employs LAMP amplification, a smartphone for fluorescence-detection and additive manufacturing as the means for scalable diagnosis in resource-limited areas. Differences to our work include the use of a disposable microfluidic unit as opposed to an Eppendorf tube (qcLAMP), more complex instrumentation (syringe pumps, filters) as opposed to a mini-camera for the colorimetric assay and a limited clinical study (10 samples) compared to ours (163 samples). A smaller, handheld device using a single plastic reaction-tube for performing RT-LAMP, a smartphone for displaying results and a sample-to-result time of <1h with an extraction-free protocol was presented in another study^65^. This fluorescence-based technology, although more complex than the qcLAMP, appears promising for the POC; verification of the methodology with more crude samples (currently 20) could provide further information on its sensitivity. Another novel isothermal amplification (RCA) assay^66^ was combined with electrochemical detection through a screen-printed electrode and a portable potentiostat. The method, validated with clinical samples and shown to provide results in less than 2 h, was using extracted RNA. Portable technologies have also been reported employing the well-established RT-PCR. The nanoPCR^67^, a device allowing fast thermocycling via plasmonic heating through magnetic nanoparticles, achieved SARS-CoV-2 detection from extracted RNA with a sensitivity and specificity of >80% and >97%, respectively and within <40 min. This sophisticated technology requires complex rotating and movable parts while its compatibility with crude samples has not been demonstrated yet. The CovidNudge platform ^68^ is the most mature POC SARS-CoV-2 device reported so far, having gone through extensive clinical validation and being already in the market. It employs a disposable lab-on-chip cartridge for swab-sample pretreatment and a 5-kilogram instrument integrating the pneumatic, thermal, imaging and mechanical parts required to run a PCR. Our methodology compares favorably to the CovidNudge due to its much simpler and more affordable technology as well as faster analysis time; on the other hand, the option for on-chip RNA purification and multiple viral-targets analysis offered by CovidNudge are clear advantages reflected in the system’s high sensitivity (95%). Finally, CRISPR-based assays have also been used for the specific SARS-CoV-2 detection employing basic infrastructure. In one such example, a smartphone was used in combination with an isothermal (RT-RPA) saliva-based COVID-19 assay with a sample-to-answer time of 15 min^69^. The end-point fluorescence detection of the assay is less accurate than real-time quantitative detection; validation results with patients’ samples will provide information on the method’s clinical sensitivity. Last, compared to a recent CRISPR-Cas-12 based method for extracted SARS-CoV-2 RNA detection via a lateral-flow assay, our method is advantageous due to its contamination-free and quantitative nature validated with crude clinical-samples; on the other hand, the double targets-detection demonstrated with the CRISPR assay could provide a more certain result ^70^. Overall, to the best of our knowledge, the swab-to-qcLAMP device and methodology reported in this work has never been shown before for SARS-CoV-2 or other viruses detection. The methodology possesses many desirable characteristics for POC diagnosis, i.e., fast analysis time, operation by minimally-trained personnel, simple protocol and instrumentation and use with commercially available reagents. It can, thus, become an attractive tool for testing in places such as airports, mobile health units, schools, nursery houses etc.

The aim of this study was to introduce a new biomedical device for performing real-time quantitative colorimetric LAMP using various samples. Based on our results, we consider two as the biggest advantages of the qcLAMP device. First, its potential compatibility with all the up-to-date reported RT-LAMP colorimetric protocols developed for both lab-based ^71^ and POC detection ^38, 59, 62, 72-74^ and for several applications in the healthcare, e.g. SARS-CoV-2 RNA detection, pathogens testing, cancer mutations etc. The second advantage is the simple and robust instrumentation, allowing fast prototyping through 3D-printing at a record time of <12 h; rapid scaling-up of the production is foreseen due to the low complexity of the design. Regarding cost, the qcLAMP prototype is comparable to a water bath typically used for heating the samples in colorimetric LAMP applications at resource-limited areas. Moreover, the handheld device is lightweight (<400 gr), can be battery-operated and is fully controlled by a smartphone. It, thus, falls well within on-going efforts to develop smartphone based POC devices for large-scale rapid testing combined with spatiotemporal surveillance and data transfer through the cloud^75^. The above characteristics make the qcLAMP device a particularly attractive solution for global diagnostics, with the potential of manufacturing and production to take place directly in the low- or middle-income countries.

So far, we have only mentioned possible applications of our device in healthcare. Nevertheless, the device can function as an open platform for performing colorimetric LAMP assays for a wide range of applications. The system could be used for the detection of food-borne pathogens at production lines with limited analytical capabilities, or for plant-borne pathogens in the field combined with on-line surveillance. In closing, we hope that this platform will be welcomed by scientists as a new tool for lab-based research but also field-measurements in a variety of applications.

## Methods

### qcLAMP device design and construction

The qcLAMP device consists of five individual parts: The main body (hosting all the electronic parts), the heating element holder, the vial holder, the bottom and top cover. All parts were made from polylactic acid (PLA) purchased from Polymaker (Netherlands) or 3DPrima (Sweden), except for the heating element holder which was made from high Tg co-polyester (HT) purchased from Colorfabb (Netherlands). HT is known for having a high glass transient temperature (∼ 100° C), thus making it a suitable material for use in hot environments. All parts were designed by using openSCAD software. The PLA parts were 3D-printed in the Bolt pro 3D printer from Leapfrog (Netherlands), while the HT parts in the TAZ6 3D printer from Lulzbot (U.S.A.). The vial holder was designed to fasten magnetically on the top cover; the latter is also connected to the main body *via* magnets. The PCB board is fixed firmly on the heating element holder, which also connects tightly to the main body. The bottom part/cover of the device encloses and protects the electronic components. The only freely moving parts of the system are the top cover and vial holder. The manufacturing protocol produces two boxes in almost 16 hours (standard speed, 0.25mm layer thickness) or 10 hours (high speed, 0.38mm layer thickness) due to the independent dual extruders that the Bolt pro offer. This way, two boxes are manufactured in parallel. The fabrication and assembly of the system can be completed within half a day.

### Electronics design and smartphone app development

The electronics layer of the qcLAMP device consists of three main components: a Raspberry Pi Zero W board (RPi), a Camera Pi module and a custom made PCB RPi Hat. The RPi operates as a central microcomputer while the Camera Pi is used for image acquisition. The PCB Hat (designed with KiCAD software) is used as an interface dedicated to control the power supply and the operation of heater, sensor and LEDs (Fig. S6). The Software layer of the device can be further divided into two categories: the software running on the RPi and the software deployed as an Android application on a mobile device. A full systems’ architecture is depicted in Fig. S7. A user-friendly interface (Android application) is used to set the temperature, colorimetric dye and an interval for the image acquisition. These initial settings are transmitted via a bluetooth low energy (BLE) connection to the corresponding bluetooth peripheral module, running inside the RPi. After setting the parameters of the experiment, a python module at the RPi end initiates the image acquisition and analysis. Each image is sent to a second python module which undertakes the image analysis procedure. The first step of this python module is to define a rectangle area on the image of the reaction tubes (Fig. S8) and continue with the application of the colorimetric analysis as described in Digital image analysis section.

### Device performance evaluation

Evaluation experiments were performed with an attenuated strain of *Salmonella enterica* serovar Typhimirium is saliva sample. Saliva from healthy donors was purchased from Lee Biosolutions, USA. A volume of 2-5μl or saliva was mixed with 1μL of *Salmonella* cells followed by cell lysis at 95 °C for 5 min, mixing with the LAMP cocktail (described in the previous section) and placing it in the device at 63°C. *Salmonella* was grown overnight in Luria−Bertani (LB) medium; cultures were subsequently measured spectrophotometrically (OD600) and adjusted at OD600:1 corresponding to a cell concentration of 1−1.5 × 10^9^ CFU/mL. Serial dilution of the above cells suspension by LB was carried out to reach the required concentration. The *Salmonella* invasion gene *invA* was targeted by a set of six primers (Metabion, Germany), two outer (F3 and B3), two inner (FIP and BIP) and two loop (Loop-F and Loop-B). The sequences of the primers (5’-3’) were as follows:

FIP:GACGACTGGTACTGATCGATAGTTTTTCAACGTTTCCTGCGG,

BIP:CCGGTGAAATTATCGCCACACAAAACCCACCGCCAGG,

F3:GGCGATATTGGTGTTTATGGGG,

B3:AACGATAAACTGGACCACGG,

Loop F:GACGAAAGAGCGTGGTAATTAAC,

Loop B:GGGCAATTCGTTATTGGCGATAG.

The LAMP reagent mix in a total volume of 25 μL contained 12.5 μL of WarmStart or Colorimetric WarmStart 2 × Master Mix (New England BioLabs) or Bsm polymerase (Thermo Scientific), 1.8 μM FIP and BIP, 0.1 μM F3 and B3, 0.4 μM Loop-F and Loop-B, and 2.5 μL of sterile water or any crude sample mixed with 1 μL of cells. The HNB color indicator was added to a final concentration of 160μM when used. LAMP was performed in the qcLAMP device at 63°C. After amplification, the products were analyzed using electrophoresis on a 2% agarose gel containing GelRed (Biotium) and visualized under UV light.

### qcLAMP testing for SARS-CoV-2

For the calibration curve, we used Influenza A as the genetic target. The primers used for Influenza A detection, designed in-house using PrimerExplorer software and purchased by Metabion (Germany), were as follows:

F3: AACAGTAACACACTCTGTCA,

B3: CATTGTCTGAATTAGATGTTTCC,

FIP: CCAAATGCAATGGGGCTACCATCTTCTGGAAGACAAGCA,

BIP: TAACATTGCTGGCTGGATCCTACAATGTAGGACCATGATCT,

LF: CCTCTTAGTTTGCATAGTTTTCCGT,

LB: CCAGAGTGTGAATCACTCTCCAC.

The DNA template used for the Influenza measurements was a PCR product of 452bp corresponding to the *HA* gene of Influenza A.

For SARS-CoV-2 detection, the primers used were targeting the *N* gene, described elsewhere^70^, were as follows:

F3: AACACAAGCTTTCGGCAG,

B3:GAAATTTGGATCTTTGTCATCC,FIP:TGCGGCCAATGTTTGTAATCAGCCAAGGAAATTTTGGGGC,

BIP: CGCATTGGCATGGAAGTCACTTTGATGGCACCTGTGTAG,

LF: TTCCTTGTCTGATTAGTTC,

LB: ACCTTCGGGAACGTGGTT.

For the clinical validation study, 2 μl of total RNA extracted from patient samples was used for RTC-LAMP. Viral RNA was extracted from 200μl of clinical samples using the NucliSens easyMAG automated system (BioMérieux, Marcy l’Etoile, France. All experiments for the calibration curve with Influenza and SARS-CoV-2 were performed with the Colorimetric LAMP mix from NEB. Real-time colorimetric LAMP (qcLAMP) was performed with our device for 30 min per run. Tests were performed using 2μl of extracted total RNA from patient samples stored at –80°C. qcLAMP was targeting a viral RNA corresponding to the N gene of the SARS-CoV-2 genome. A second RNA target (*ORF1a*) was also available but selectively tested (data not shown). A human RNA target (*RNase* P) was used as a positive reference with a previously described set of primers^70^. No-template reactions were periodically performed for checking the quality of the reagents (e.g. contamination) and the temperature stability of the device. Synthetic SARS-CoV-2 RNA was purchased from BIORAD (SARS-CoV-2 Standard #COV019 and SARS-CoV-2 Negative #COV000).

For the direct detection of SARS-CoV-2, 100 μL of each clinical sample have been diluted 1:1 in neutralizing buffer and remained at RT for few minutes. 5 μL of the dilution was added to the LAMP reagent mix in a total volume of 25 μL. The mix contained in final concentration 8U *Bst DNA/RNA Polymerase* from SBS Genetech Co. (Beijing SBS Genetech Co., Ltd.), 10× Bst DNA/RNA polymerase isothermal buffer (Mg^2+^ free), 7 mM Mg^2+^, 1.2 mM (each) dNTPs (Thermoscientific), 160 μM HNB dye and nuclease-free water up to 25 μL. The primers used were targeting the *N* gene described in the previous section. The qcLAMP reaction was performed in the device at 63°C for 35 min.

### Sample collection for SARS-CoV-2 testing

Nasopharyngeal or oropharyngeal swabs were acquired from patients with suspicion of COVID-19 infection. The Institutional review board (IRB) guidelines were followed. Clinical swab samples were collected in VTM from several public and private hospitals of Greece and forwarded to the two National SARS-CoV-2 reference laboratories participating in this study: the Hellenic Pasteur Institute (Athens, Greece) and the Laboratory of Clinical Virology at the School of Medicine, University of Crete, Greece.

### qRT-PCR assays for SARS-CoV-2

#### Hellenic Pasteur Institute

Viral RNA was extracted from 200μl of clinical samples using the NucliSens easyMAG automated system (BioMérieux, Marcy l’Etoile, France). A 106bp fragment of the SARS-CoV-2 virus *RdRp* gene was amplified according to an in-house qPCR protocol proposed by the WHO and the National Reference Center for Respiratory Viruses, Institut Pasteur, Paris. As a confirmatory assay, the *E* gene assay from the Charité protocol was used^76^ by real-time RT-PCR. SeraCare’s SARS-CoV-2 AccuPlex solution 5000copies/ml (Material number 0505-0126) was used for limit-of detection testing. 1 μL of the solution was directly placed in the LAMP mix or after 10 min of heating at 80°C.

#### Laboratory of Clinical Virology, Un. of Crete

Viral RNA was extracted from 200μl of VTM using the MagMAX^™^ Viral/Pathogen II (MVP II) Nucleic Acid Isolation Kit (Applied Biosystems) on the KingFisher^™^ Flex Purification System (Thermo Scientific) according to the manufacturer’s protocol. QPCR COVID-19 detection was perfomed using the TaqPath COVID-19 CE-IVD RT-PCR kit (Applied Biosystems) on the QuantStudio 5 real-time PCR (Applied Biosystems) with the associated Applied Biosystems COVID-19 Interpretive Software. The TaqPath COVID-19 RT-PCR kit targets genomic fragments in the ORF gene, *S* gene and the *N* gene. It uses internal control for assessing extraction efficiency and PCR inhibitors.

### Detection of *BRAF* V600E mutation from extracted genomic DNA

Loop-mediated amplification was conducted by using genomic DNA, *BRAF* V600E Reference Standard and *BRAF* Wild Type Reference Standard, purchased from Horizon Discovery or extracted from formalin-fixed and paraffin-embedded (FFPE) tissue (clinical samples). A set of six primers (Metabion, Germany) were used for the detection of *BRAF* V600E mutation; two outer (F3 and B3), two inner (FIP and BIP) and two loop (Loop-F and Loop-B). The sequences of the primers (5’-3’) were as follows:

FIP:TCTGTAGCTAGCAGATATATTTCTTCATGAAGACCT, BIP: AGAAATCTCGATTCCACAAAATGGATCCAGA, F3: GGAAAATGAGATCTACTG, B3: TCTCAGGGCCAA, LF: ACCAAAATCACCTATTT, LB: GGAGTGGGTCCC.

LAMP was carried out in a total of 25μl reaction mixture containing 1.6μM of the forward inner primer (FIP), 1.6μM of the backward inner primer (BIP), 0.2μM of the forward outer primer (F3), 0.2μM of the backward inner primer (B3) and for the loop primers 0.8μM of loop-forward (LF) and 0.8μM of loop-backward (LB) as described previously^29^. The total concentration of genomic DNA used was 100ng/μl, which contained wild-type and/or mutant DNA, adjusted depending on the number of mutant copies to be detected (50%-0.01% mutant DNA). Positive and negative controls were included in each set of reactions and every precaution needed in order to avoid cross-contamination. 2 μl of genomic DNA were mixed with LAMP primers, denatured for 5 min at 92°C and quickly placed on ice. The solution was subsequently mixed with 12.5 μL WarmStart® colorimetric LAMP and the reactions were finally placed in the qcLAMP device for real-time amplification monitoring at 63°C.

### Direct detection of *BRAF* V600E from formalin fixed paraffin tissue

Tissue slices of various mutant:wild type ratios for the *BRAF* V600E mutation were purchased from Horizon Discovery. Slices of 15 μm (or 1 mm in length) were immersed in 100 μl of ddH_2_O and heated at 95°C for 65 min followed by a brief centrifugation step at 13000 rpm for 2 min. 10 μl of the solution were then mixed with the LAMP primers and denatured at 92°C for 5 min. For the 1% ratio, a 100% wild type solution was mixed with a 10% to the final volume of 10 μl. After the denaturation step, each solution was placed on ice before mixing it with 12.5 μL of WarmStart® colorimetric LAMP and placing in the bioPix device for real-time amplification monitoring at 63°C.

### Formalin-fixed and paraffin-embedded (FFPE) tissue processing and DNA extraction of the clinical samples

Twelve FFPE samples from patients with cancer were investigated, including 4 melanomas, 4 colorectal cancer tissues and 4 non-small cell lung cancer samples. The research protocol was approved by the Ethics Committee of the University Hospital of Heraklionand all participants provided written informed consent. Tumor areas were micro-dissected to enrich the analyzed specimen with cancer cells and DNA was extracted using the QIAamp DNA Micro Kit (Qiagen). DNA concentration was determined spectrophotometrically by the absorbance measurement at 260nm and all samples were kept at −20oC until use.

### Sanger sequencing for the clinical samples

Samples from cancer patients were tested by Sanger sequencing, which is a gold standard method for detecting DNA mutations. In more details, a PCR assay was designed to amplify *BRAF* exon 15, which includes the mutation hotspot that encodes the V600E variant. The primer sequences used in this study were as follows: *BRAF* forward 5’-TGT TTT CCT TTA CTT ACT ACA CCT CA-3’ and reverse 5’-GCC TCA ATT CTT ACC ATC CA-3’ (PCR amplicon of 160bp). PCR products were then electrophoresed on 2.0 % (w/v) agarose gel to check the presence of specific amplification products. Unincorporated PCR primers and dNTPs were removed from the PCR products using the Nucleospin PCR clean-up kit (Macherey-Nagel) according to the manufacturer’s protocol. The sequencing reactions were performed in a final volume of 10.0 μl using: 2.0 μl of purified PCR product, BigDye Terminator v3.1 Sequencing reagents (Applied Biosystems), and 125nM each of the forward and reverse primers. The products were purified using ethanol precipitation and Sanger sequencing was performed by capillary electrophoresis (ABI3130, Applied Biosystems). Sequencing analysis v5.4 software (Applied Biosystems) was used to analyze the results.

### Droplet Digital PCR for *BRAF* V600E Quantification

DNA samples with the *BRAF* V600E mutation, as determined with Sanger sequencing, were by the ddPCR system for absolute quantification of mutant alleles and wild-type alleles. The ddPCR was performed using the QX200 Droplet Digital PCR System (Bio-Rad) and the ddPCR *BRAF* V600 Screening Multiplex Kit (Bio-Rad) for 3 different hot-spot codon V600 mutations (V600E, V600K, and V600R), according to the manufacturer’s protocol. Each sample was tested in two technical replicates and every ddPCR run included negative template controls (NTCs) and positive controls (*BRAF* mutant homozygous and heterozygous cancer cell lines and Wild-Type (WT) samples). Samples were mixed with 70.0μL of droplet generator oil for probes (Bio-Rad) and partitioned into up to 20,000 droplets using the Bio-Rad QX-200 droplet generator (Bio-Rad). Then, 40.0μL of emulsion for each sample, was transferred to 96-well plates (Bio-Rad) and PCR was done on C1000 Touch thermal cycler (Bio-Rad), using the following conditions: 95°C for 10 min, and 40 cycles of 94°C for 30 s, 55°C for 1 min, and 98°C for 10 min. The plate was then transferred and read in the FAM and HEX channels using the QX200 droplet reader (BioRad). Data analysis was performed using the QuantaSoft Analysis Pro Software (Version 1.0.596) to assign positive/negative droplets and convert counts to target copies/μL of reaction. Threshold was manually set for each sample, based on positive control samples (mutant and WT for each channel).

## Data availability

The authors declare that all data supporting the findings in this study are available within the paper and its Supplementary Information files.

## Funding

This work has received funding from the EC through the H2020-SC1-PHE-CORONAVIRUS-2020-2B grant No 101016083 (project acronym “IRIS-COV”), H2020-FET-OPEN-2018-2019-2020-01 grant No 862840 (“FREE@POC”), H2020-FETOPEN-1-2016-2017 grant No 737212 (“CATCH-U-DNA”) and the Patras Science Park “Proof-of-Concept” grant No: 1718B (“BioPix”).

## Author Contributions

G.P. conceived the real-time colorimetric LAMP idea and digital image analysis concept, performed the proof-of-concept and evaluation experiments, participated in the design evolution of the device and analyzed all data. A.K.P. designed, manufactured and assembled the device; assisted with the proof-of-concept evaluation, the HAT development and digital image analysis and co-designed the heaters together with N.F. N.F. developed the HAT-unit and the smartphone application and participated in the design evolution of the device. M.M and M.V. carried out all performance evaluation experiments including qcLAMP *Salmonella* detection. J.H. and E.N. prepared and provided reagents (Influenza A DNA). S.C. and V.P performed qcLAMP (Influenza, SARS-CoV-2) and qRT-PCR (SARS-CoV-2) experiments. V.T. performed all qcLAMP experiments using crude patients sample. K.G. carried out the qcLAMP *BRAF* V600E detection. T.K. and A.M. supervised clinical evaluation experiments for COVID-19 at the Hellenic Pasteur Institute A.Z and G.S. at the Laboratory of Clinical Virology at the Univ. of Crete. S.A and K.M. performed Sanger sequencing and ddPCR experiments for *BRAF* mutation detection and analyzed the data. E.G. supervised the whole work and together with G.P. wrote the manuscript. All authors read, contributed to the manuscript and agreed to its content.

## Competing interests

G.P., A.K.P. and E.G. are inventors of one patent related to this work filed by FORTH (PCT/EP2019/079845). G.P., A.K.P., N.F. and E.G. are the co-founders of BIOPIX DNA TECHNOLOGY P.C., a spin off company exploiting the current technology and relevant patent. The other authors declare no competing interests.

## Data and materials availability

All data needed to evaluate the conclusions in the paper are present in the paper and/or the Supplementary Materials. Additional data may be requested directly from the authors.

## Notes

### Competing Interest Statement

G.P., A.K.P., N.F. and E.G. are the co-founders of BIOPIX DNA TECHNOLOGY P.C. The other authors declare no competing interests

### Summary of Updates

To fully exploit the qcLAMP methodology, we also developed a swab-toqcLAMP assay combined with reverse-transcription, circumventing the RNA-extraction step and, thus, suitable for POC testing.

