## Supplementary Material for "Portable real-time colorimetric LAMP-device for rapid quantitative detection of nucleic acids in crude samples"

#### In-house developed Android application

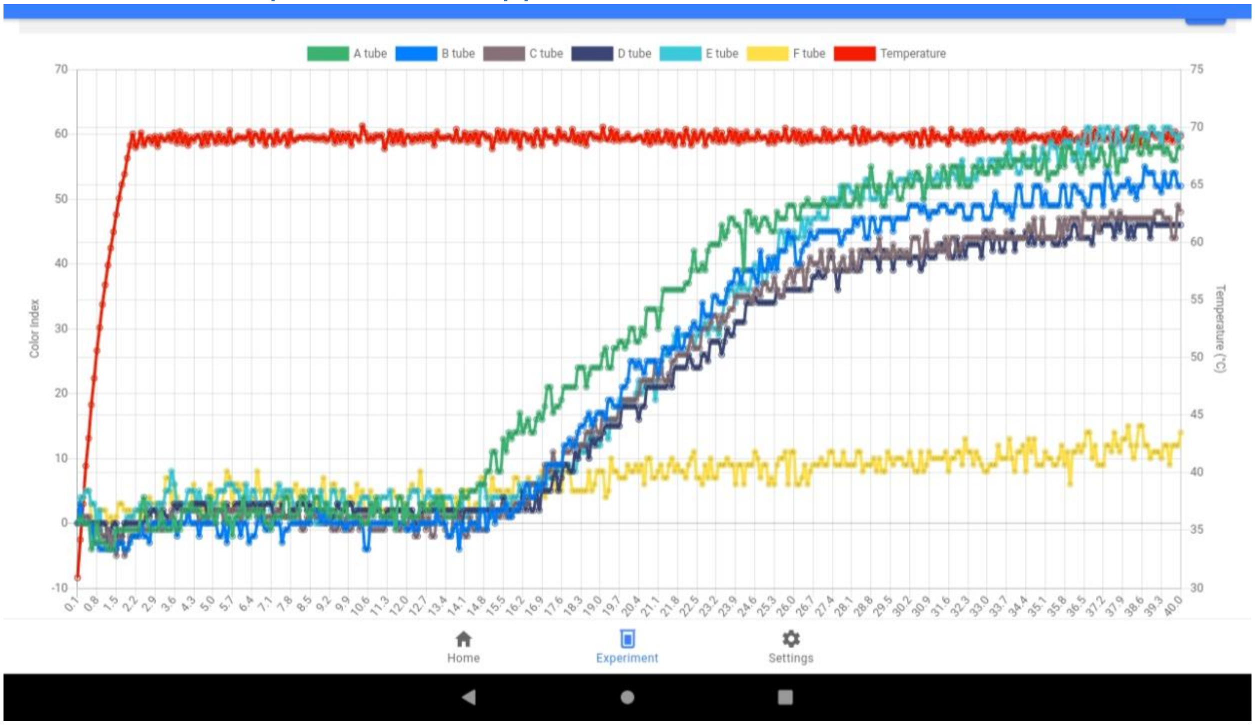

**Fig. S1**: Screenshot of the in-house developed Android application. The settings to be adjusted include the temperature, run time, type of dye, time interval for capturing images and option for USB storage.

### Digital image analysis

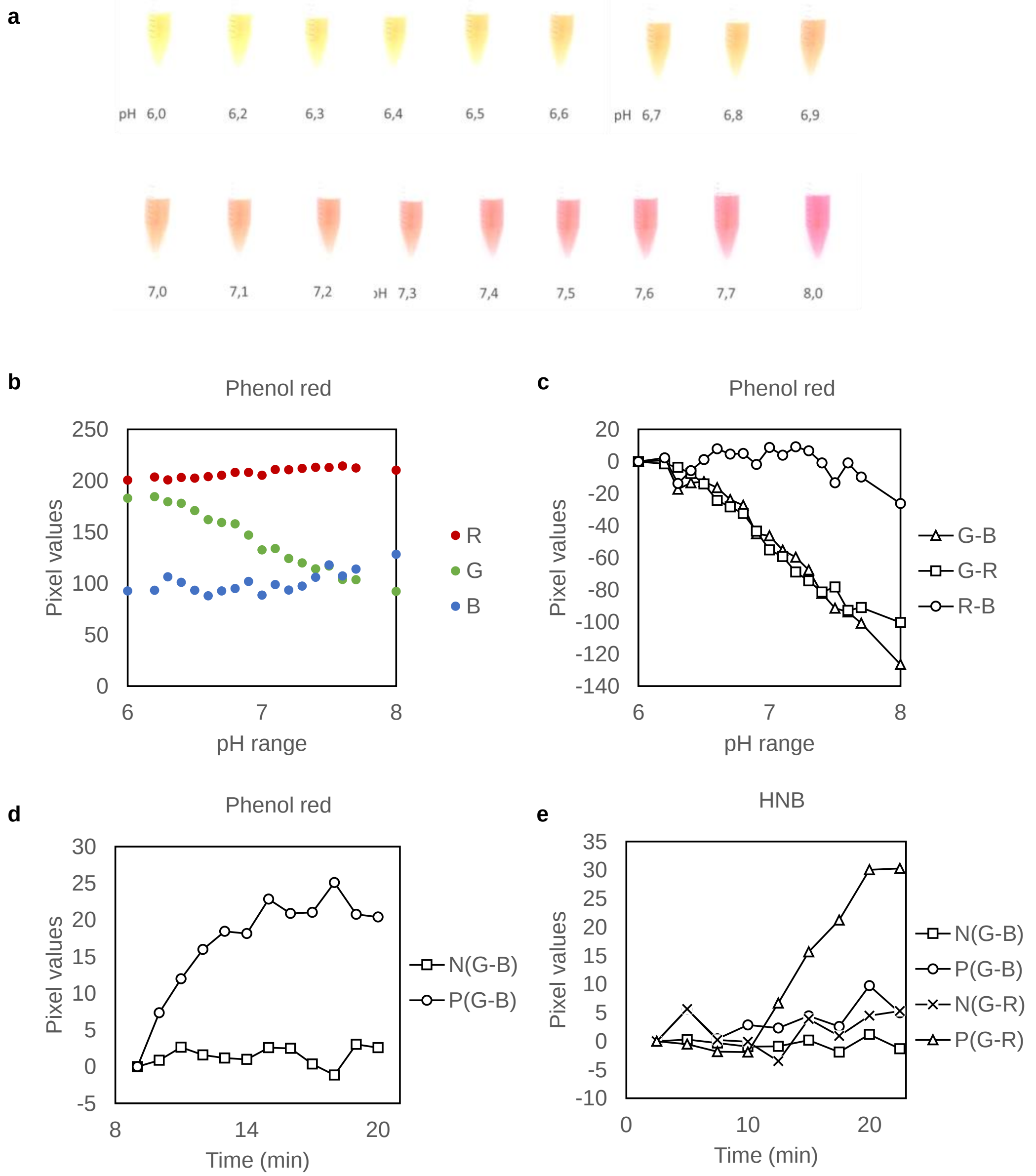

**Fig. S2**: **Digital image analysis.** (a) Series of images corresponding to different pH values between pH 6 and 8 in respect to the phenol red indicator. Image source: https://www.testallcolour.com/blog/post/what-is-phenol-red-in-swimming-pools/. (b) Raw pixel values extracted from several images (see a) correlating different pH values to color change using the phenol red pH indicator. (c) phenol red: change in pixels as function of pH using data from figure S2b and following three formulas; Green-Blue (G-B), Green-Red (G-R) and Red-Blue (R-B). (d) Change in pixels of phenol red based LAMP reactions spiked with 0 (N) and $10^5$ (P) lysed bacteria by applying the Green-Blue formula. The first 8 minutes were omitted. Reactions took place in a pre-warmed oven at 63°C with a glass door that allowed video capturing with a camera placed outside the door. (e) Change in pixels of HNB based LAMP reactions spiked with 0 (N) and $10^5$ (P) bacteria by applying the Green-Blue and Green-Red formulas. With the HNB indicator, the Green-Red formula resulted in better discrimination (Fig. S2e); this could be explained by the fact that the purple to sky blue transition involved more prominent changes in the green and red than in the blue channel.

### Performance evaluation

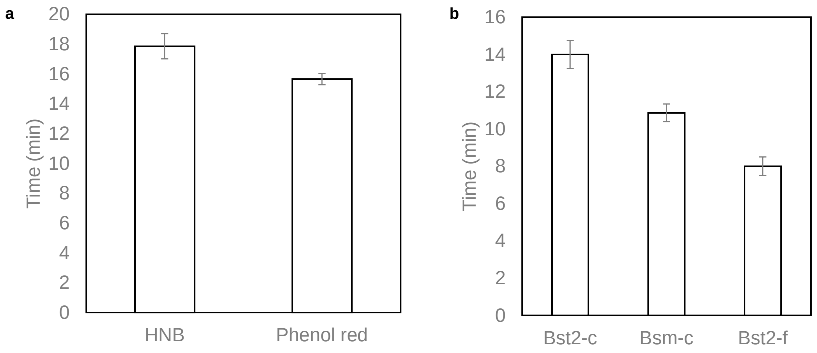

**Fig. S3**: **LAMP speed of detection.** (a) Variation in the time point (min) in which LAMP preparations (containing the same amount of starting template but different color indicator) show a change in the slope of the real-time colorimetric curve when changing the position of the tube inside the tubes holder (see Fig. 1b). Each bar is the average of 3 replicates at 2 different slots in the holder (total of 6 measurements). (b) Comparison of the speed of detection of a LAMP reaction containing 10 bacteria as starting template using different combinations of 2 enzymes (Bst2, Bsm), 2 colorimetric indicators (HNB, phenol red) and inside 2 real time systems (qcLAMP device, BIORAD). Bst2-c: Bst2 warm start polymerase mixed with either phenol red or HNB, tested with qcLAMP; Bsm-c: Bsm polymerase (20 Units) with HNB, tested with qcLAMP; Bst2-f: Bst2 warm start with LAMP fluorescent dye tested in a real-time PCR machine. Error bars represent standard deviation of at least triplicate measurements.

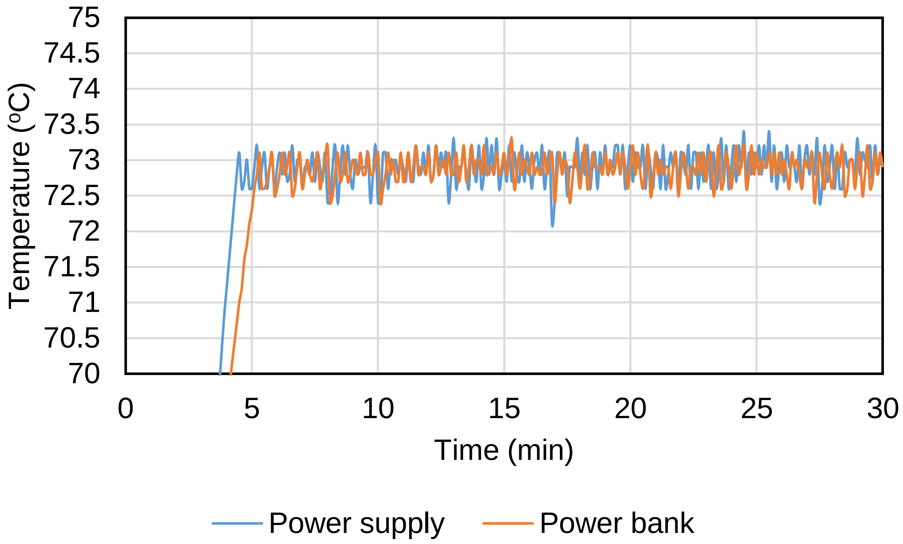

**Fig. S4**: **Temperature stability of the device.** Temperature stability during operation of the qcLAMP device with a power bank and in comparison, to a standard power supply.

#### qcLAMP curves for Influenza/SARS-CoV-2

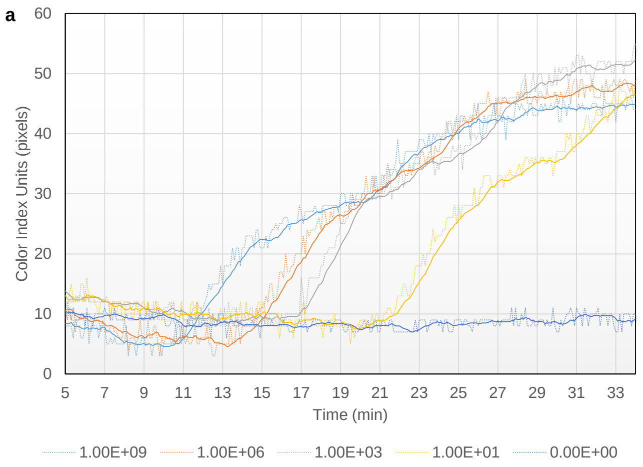

b

| NUMBER OF SAMPLE | TIME-TO-POSITIVE (MIN) |
|---|---|
| 1 | 9.7 |
| 2 | 10.2 |
| 3 | 10.9 |
| 4 | 10.1 |
| 5 | 11.7 |
| 6 | 10.2 |
| 7 | 9.4 |
| 8 | 10.5 |
| 9 | 10.4 |
| 10 | 10.1 |
| 11 | 9.2 |
| 12 | 10.6 |
| 13 | 10.6 |
| 14 | 10.2 |
| 15 | 9.9 |
| 16 | 10.2 |
| 17 | 9.6 |
| 18 | 9.9 |
| 19 | 9.7 |
| 20 | 10.5 |
| 21 | 10.7 |
| AVARAGE TIME (MIN) | 10.2 |
| STANDARD DEVIATION (min) | 0.6 |

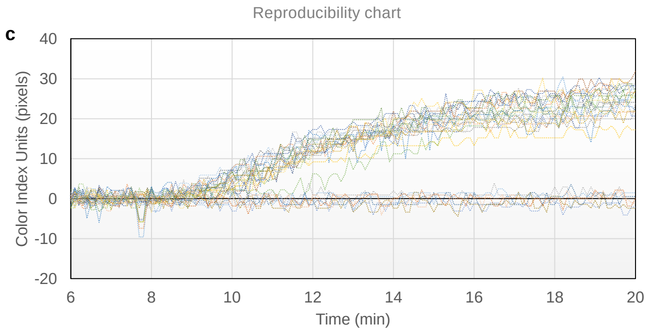

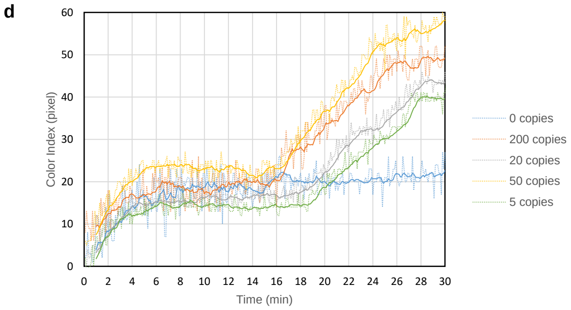

**Fig S5**: **qcLAMP tests with Influenza A and SARS-CoV-2 template.** (a) Typical real time colorimetric LAMP curves for Influenza A. (b) Average time-to-positive for 21 positive samples with the same initial target concentration ($10^9$ copies/reaction). (c) Real-time curves of 28 samples (21 positive, 7 negative). (d) Real time curves for 0 to 200 copies of SARS-CoV-2 synthetic RNA.

#### Electronics design and smartphone app development

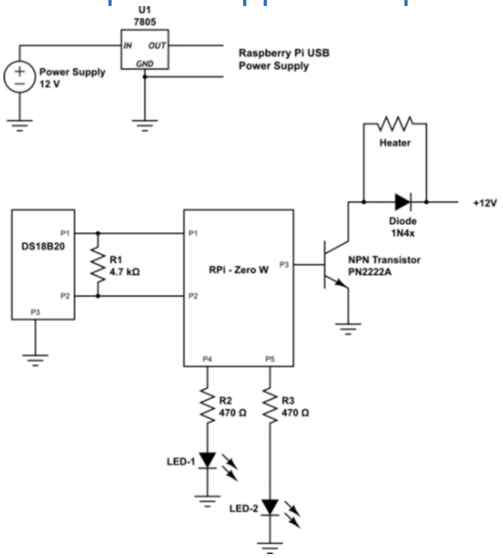

**Fig. S6**: **Electronics design.** Schematic representation of the custom PCB RPi Zero W Hat for controlling temperature sensor (DS18B20), Heating element, and LEDs.

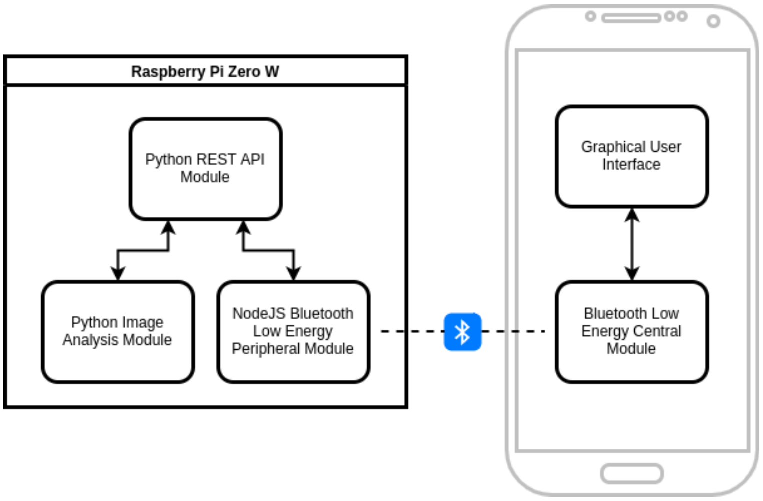

**Fig. S7**: **Systems' architecture at software layer.** The Software layer of the device is divided into two categories: the software running on the RPi and the software deployed as an Android application on a mobile device.

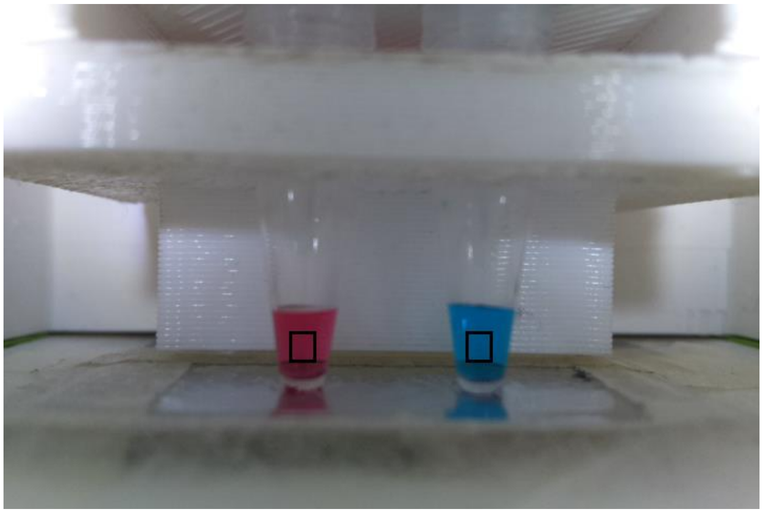

**Fig. S8**: **Snapshot of the reaction tubes in the qcLAMP device.** The application of the colorimetric analysis is performed in predefined areas depicted in the black rectangles. Photo Credit: N. Fikas Photographer Institution: IMBB-FORTH.
